# Fast feature- and category-related parafoveal previewing support natural visual exploration

**DOI:** 10.1101/2024.04.29.591663

**Authors:** Camille Fakche, Clayton Hickey, Ole Jensen

## Abstract

Studies on vision tend to prevent or control eye movements, while humans naturally saccade every ∼250 ms. As the oculomotor system takes ∼100 ms to initiate and execute a saccade, this leaves only ∼150 ms to identify the fixated object and select the next saccade goal. This is very little time, suggesting that vision relies on parafoveal processing before and after the eye movement. However, evidence of high-level parafoveal access is sparse. The purpose of our study was to use magnetoencephalography (MEG) combined with eye-tracking and multivariate pattern analysis to identify the neuronal dynamics of parafoveal processing which support natural visual exploration. We demonstrated that future saccade goals in the parafovea could be decoded at the feature and category level peaking at ∼90 ms and ∼160 ms respectively. Simultaneously, decoding of fixated objects at the feature and category level peaked at ∼70 ms and ∼145 ms respectively. Also decoding feature and category specific neuronal information related to past parafoveal objects were sustained for ∼230 ms after saccading away from them. The feature and category of objects in the parafovea could only be decoded if they were in the saccade goal. In sum, we provide insight on the neuronal mechanism of pre-saccadic attention by demonstrating that feature and category specific information of foveal and parafoveal objects can be extracted in succession within a ∼150 ms time-interval and may serve to plan the next saccade. This information is maintained also after fixations and may support integration across the full visual scene. Our study provides novel insight on the temporal dynamics of foveal and parafoveal processing at the feature and semantic levels during natural visual exploration.

## Introduction

Humans have a remarkable ability to explore visual scenes efficiently. This capacity relies on eye movements occurring every ∼250 ms, which shifts the foveal view to informative parts of the visual scene [1, 2]. Each inter-saccade interval includes parafoveal processing (2 – 5° of eccentricity) and has three likely functions. The first function is to support the preparation of saccade goals. Since our eyes typically land on informative parts in a visual scene, a selection must be made between candidates for saccade goals. The second function serves to give visual processing a head-start, i.e., the parafoveal previewing of an object serves to speed up the processing of the object when fixated. The third function includes post-saccadic memory. As a visual scene is perceived as stable despite frequent eye-movements, information preceding each saccade must be integrated with visual input following the saccade. Since we saccade every ∼250 ms, and the oculomotor system takes 100 ms to initiate and execute a saccade, the visual system has roughly 150 ms to complete these three functions [3]. The purpose of our study is to use magnetoencephalography (MEG) combined with eye-tracking and multivariate pattern analysis to identify the neuronal dynamics of parafoveal processing which support natural visual exploration. Specifically, we seek to identify the time course of parafoveal processing and determine the kind of information is resolved.

Recent studies have argued that parafoveal target goals are processes even at the semantic level. Previous studies found that objects incongruent with a given scene were fixated more often and for a longer duration than congruent objects [4–6]. Objects semantically incongruent within the scene were fixated earlier than congruent ones. The probability of first fixation was also higher for semantically-incongruent objects [4–7]. These effects are best explained by target goals being processed even prior to a saccade. In further support, a visual search task demonstrated that target objects were fixated earlier and more often when they were semantically unrelated to the distractors [8, 9]. These findings provide circumstantial evidence for semantic parafoveal processing guiding saccades during visual search. One electroencephalography (EEG) study has provided electrophysiological evidence for contextual visual information being derived parafoveally during natural scene viewing [10]. In a free-viewing task, the congruency of a parafoveal visual object within a scene, indexed by the N400, was measured during foveation. However, the effect was observed at ∼400 ms, so it was too slow to reflect the preview benefit and to impact the next saccade plan. Therefore, whether objects in the parafovea can be processed sufficiently fast semantically to impact saccade goals in natural viewing conditions remains an open question. The first aim of our study was to investigate whether a parafoveal previewing of visual objects at the semantic level was identifiable in the brain data in the intersaccadic interval.

A key purpose of parafoveal processing might be to reduce the processing time when the object is fixated after a saccade, an effect termed preview benefit or preview positivity [11, 12]. The preview benefit has been studied using gaze-contingent paradigms. In such paradigms, participants are instructed to fixate and then saccade towards a target object, presented parafoveally, that either changes or remains the same during the saccade. Results show that the naming of parafoveal objects is faster (∼30 to ∼150 ms) when they were previewed [13–15]. The preview benefit is further mediated by the spatial proximity of fixated and parafoveal objects, with closer stimuli benefitting more from preview [13–15]. Similarly, fixations on objects were reduced by ∼100 ms when objects were previewed [16]. Previewing an object also allows for improved detection of low- and high-level visual features [17–25]. Electrophysiological studies have found that the amplitude of the visual evoked potentials (ERP) elicited by an object was reduced after the object was parafoveally previewed as compared to not previewed [24–27]. This preview effect dramatically decreased when noise was added to the parafoveal stimulus for faces and objects [25, 27]. Another ERP study revealed that the onset latency of the N170 illicit by a face was shortened if the face had been previewed [24]. Multivariate pattern analysis applied to EEG data showed that the peak of classification accuracy associated with the categorization of foveal objects emerged ∼30 ms earlier (from 150 to 120 ms) when previewed [28]. In sum, these studies provide evidence for parafoveal previewing at the semantic level and show that previewing can reduce the processing time of an object when fixated. Nevertheless, these studies employed paradigms in which saccades either were prevented or controlled. We here complement these findings by investigating parafoveal previewing allowing natural visual exploration. In particular, we aimed to compare the time course of parafoveal and foveal visual information processing.

Visual scenes are perceived as stable despite frequent eye-movements. The stable perception may be supported by trans-saccadic memory from one fixation to the next [29–31]. Some studies have found that human participants combine the parafoveal and the post-saccadic foveal view using a weighted sum, thus providing evidence for trans-saccadic memory [19, 32]. A recent study found trans-saccadic perceptual fusion of two different stimuli: Participants were presented with a vertical line before the saccade, and with postsaccadic horizontal lines, and they perceived a fusion of the two lines in 67% of trials. The fused percept provides evidence for trans-saccadic memory of the previously viewed object [33]. Using electrophysiological approaches, multivariate classification analysis has been used to investigate whether the visual features of presaccadic stimulus could be classified after saccades. This work has shown that the spatial frequency of a presaccadic stimulus could be decoded from the EEG data ∼200 ms after the saccade [34]. The study further reported a 100 ms interval after fixation where both presaccadic and postsaccadic spatial frequencies could be decoded [34]. Similarly, another EEG study found that the category of a presaccadic object could be classified ∼100 ms after the saccade. Interestingly, this information disappeared in the EEG data when no saccade was performed [28]. In sum, there is both behavioural and electrophysiological evidence supporting trans-saccadic visual memory. These findings on trans-saccadic memory have been obtained using highly operationalized paradigms including gaze-contingent visual displays and cue-guided saccades. The third aim of our study was to complement the findings on trans-saccadic memory using a paradigm allowing free visual exploration. In particular, we set out to test whether presaccadic visual features were reflected in the multivariate brain data after saccades.

Here, we investigated the neuronal dynamics associated with feature- and category-related processing of parafoveal objects during natural viewing. Specifically, we quantified how fast and in which detail parafoveal objects are processed before the saccade onset. We also identified the time course associated with the classification of presaccadic, foveal, and postsaccadic visual features. To this end, we designed a free-viewing task using a set of natural images, displayed either in grey-or colour scale, and belonging to one of three different categories (animal, food, object). Participants were encouraged to freely explore the images while eye-tracking and the MEG data were simultaneously recorded. Multivariate pattern analysis was used to classify when the feature (colour) and the category of past, fixated, and upcoming parafoveal images can be decoded. Our main finding was that the feature and category of foveal images and parafoveal saccade goals can be identified in succession from the MEG data within ∼150 ms, providing evidence for semantic parafoveal previewing.

## Results

Participants were asked to freely explore 7 natural images using eye movements in 4 s long trials (**Figure 1A**). The images could be in colour or greyscale and belong to one of the three categories: animal, food, or object. These images were selected from the THINGS database [35]. The display was then masked for 2 s, and then 6 of the images were presented again while one was changed (a different image from the same category and colour category). The task of the participants was to identify the changed image by button press. The aim of the task was to encourage participants to explore the 7 images during the initial presentation.

**Figure 1.**
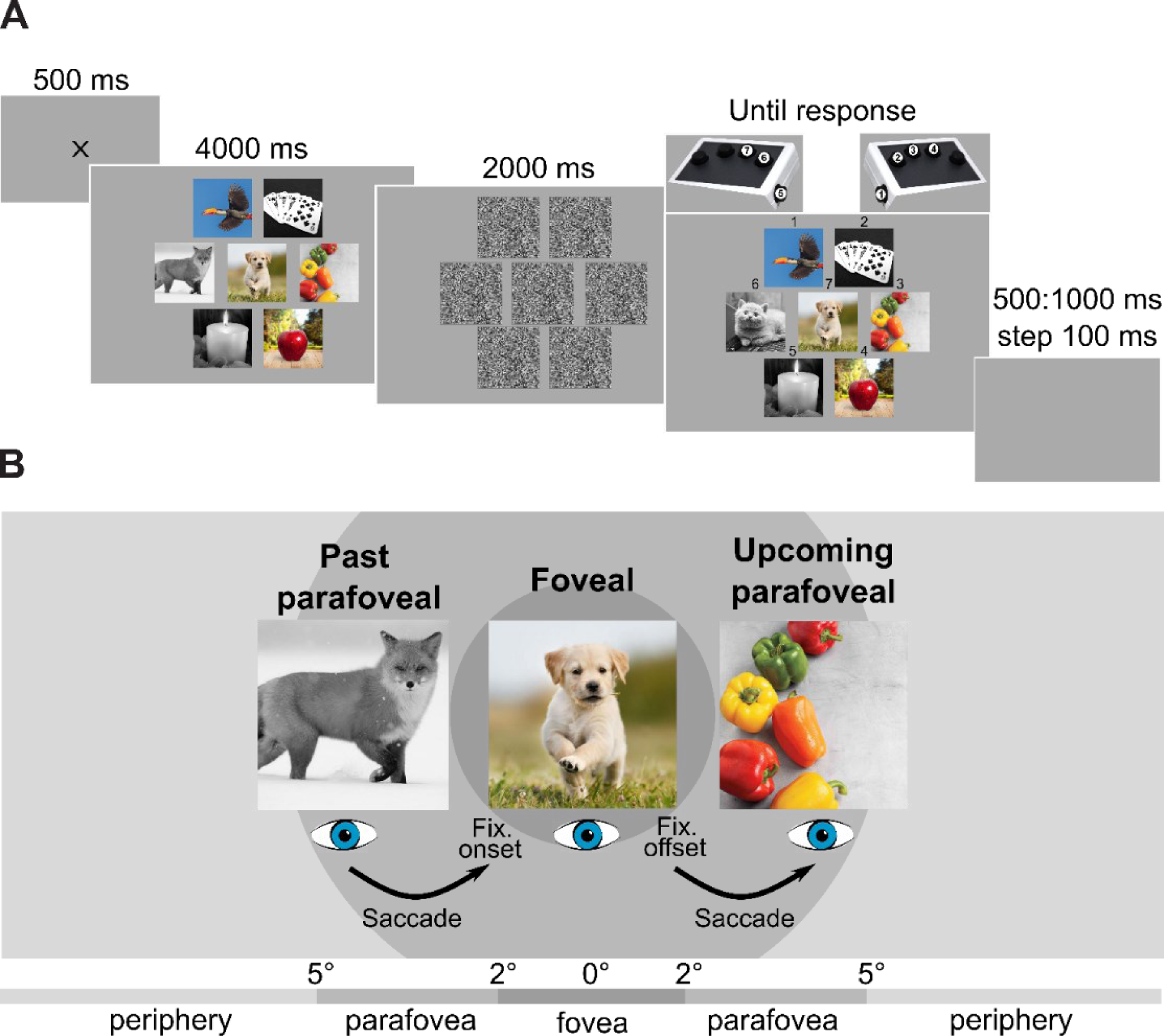
Visual exploration of natural images by saccades. **A.** Each trial started with the presentation of a central fixation cross for 500 ms. Then followed 7 natural images displayed for 4000 ms. Natural images were shown using either a grey or colour scale, and they belong to one of the three different categories: animal, food, or object. The proportion of colour and categories were balanced within and between trials. Participants were asked to freely explore the images and eye-movements were allowed. Then followed a mask for 2000 ms, after which the 7 images were presented again. One of the images had changed: the different image, belonging to the same category and colourscale as the initial one. In this example, the greyscale fox turned into a greyscale cat (6th position). The participants had to identify, without time-limit, the image that had changed. A number was presented above each image, as well an imaging showing the link between the image numbers and the response buttons. Each trial was interleaved with a 500 – 1000 ms random delay. **B.** The processing of the colour and the category of natural images was investigated in three conditions: 1) The fixated image in the fovea. 2) Upcoming image in the parafovea, corresponding to the image that will be fixated after the saccade. 3) Past image in the parafovea, corresponding to the image that was viewed before the current fixation.

### Behavioral and Eye data

All participants were able to identify the changed image as reflected by a hit rate being all above chance (67.7 ± 12.3 %; mean ± std; **Figure 2A**). As expected, the median reaction time was longer for incorrect (4002 ± 991 ms) compared to correct (2810 ± 628 ms) responses (one-tailed t-test: t(52.45) = −5.66, p < 0.001, Cohen’s d = 1.41, CI95% = [-inf, −0.84**]; Figure 2B**).

**Figure 2.**
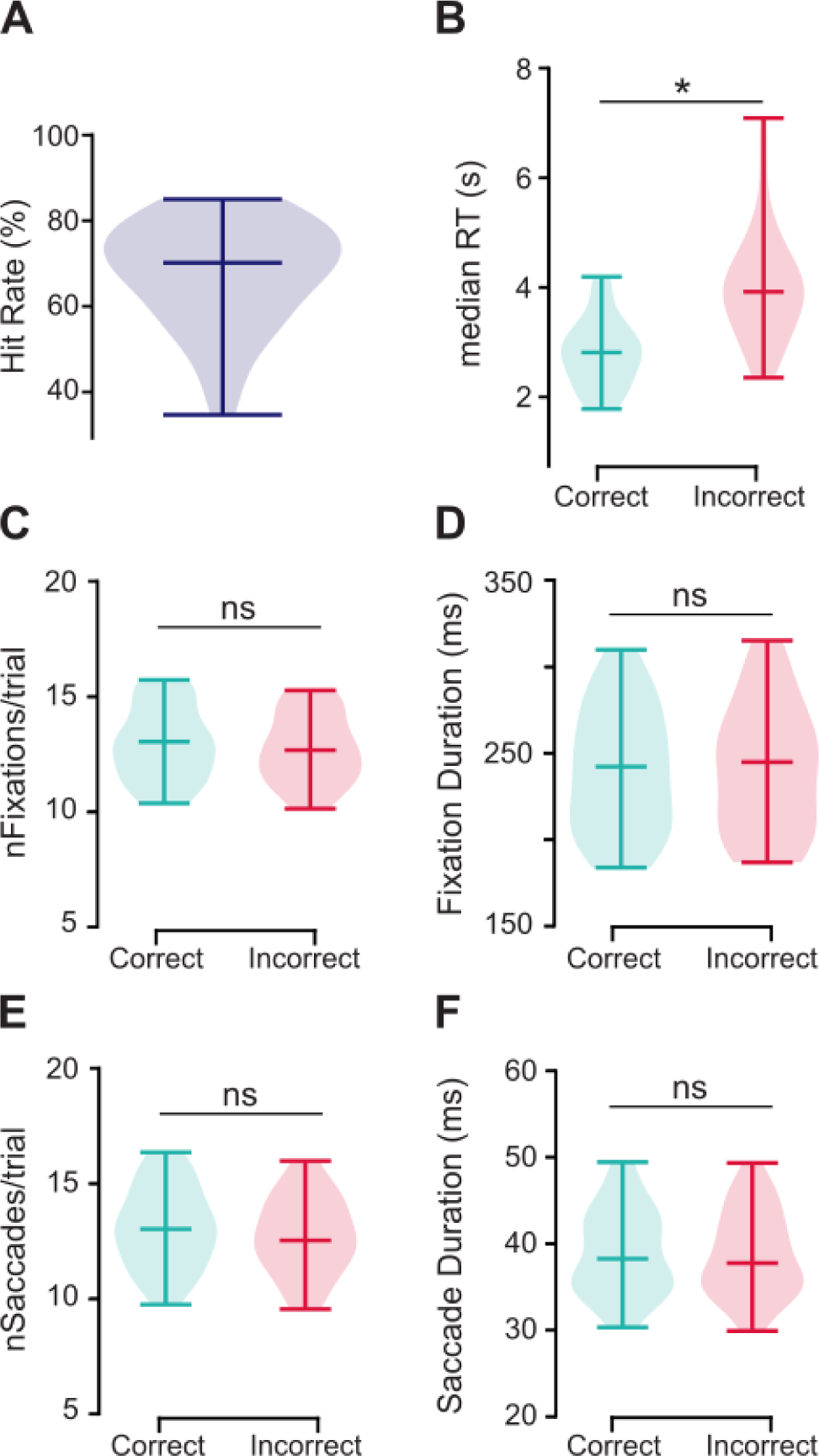
The eye metrics did not predict behavioural performance. **A.** Hit rate regarding identifying the image that had changed, **B.** The median reaction times were significantly longer for incorrect compared to correct trials (one-tailed t-test: t(52.45) = −5.66, p < 0.001, Cohen’s d = 1.41, CI95% = [-inf, −0.84]). There was no significant difference between correct and incorrect trials for the eye movement metrics (C, D, E, and F). **C.** The number of fixations per trial for correct and incorrect trials. **D.** The average fixation durations for on each object, **E.** Number of saccades per trial for correct and incorrect responses, **F.** The mean saccade duration (i.e., eyes in flight). The horizontal bar in the violin plots indicates the mean value. Dark blue plot, all trials. Cyan plots, correct trials. Pink plots, incorrect trials. ns, non-significant. *, p < 0.001.

Interestingly, the eye movement metrics did not reflect the behavioural performance. There was no significant difference between correct and incorrect trials in regard to the number of fixations per trial (two-tailed t-test: t(61.72) = 0.61, p = 0.54, Cohen’s d = 0.15, CI95% = [−0.54, 1.02], **Figure 2C**), the fixation durations were similar (two-tailed t-test: t(61.92) = −0.096, p = 0.92, Cohen’s d = 0.024, CI95% = [−0.02, 0.02], **Figure 2D**), the number of saccades per trial was similar (two-tailed t-test: t(61.92) = 0.83, p = 0.41, Cohen’s d = 0.21, CI95% = [−0.51, 1.23], **Figure 2E**), and the saccade duration was similar (two-tailed t-test: t(61.97) = 0.17, p = 0.86, Cohen’s d = 0.043, 95%CI = [−0, 0]) (**Figure 2F**). We conclude that eye movement metrics were similar for correct and incorrect trials. Participants did on average 3.23 ± 0.43 saccades per s, in line with the previous literature [36]. The mean fixation duration on each of the natural images was 240.6 ± 36.0 ms.

### MEG data

Multivariate pattern analysis (MVPA) was applied to the MEG data aligned according to the fixation onset on foveal images to investigate whether we can classify the colour (greyscale vs colours) and the category (animal vs food vs object) of the:

- foveal images (currently fixated images),
- parafoveal upcoming images (viewed after the current image),
- parafoveal past images (viewed before the fixation on the current image),
- one parafoveal remaining image (not viewed after, not viewed before the current image), as illustrated in **Figure 1B**.

### Foveal decoding features and categories

For foveal fixations, the classifier was trained and tested at each time point (using a 50 ms sliding time window and 5-fold cross-validation). To reduce contamination by the upcoming or previous saccades, we focused our interpretation of the classifier results in the −250 – 250 ms interval aligned to the fixation onset. As seen in **Figure 3A** (blue curve), the classifier could reliably distinguish the colour of the foveal images well above chance level (AUC of 0.5) in the 8 – 310 ms interval (p < 0.05; cluster-permutation approach controlling for multiple comparisons over time [37–39]). Classification performance (AUC) gradually built up until peaking to 0.69 at 70 ms, after which they began to decrease. The classifier could also reliably distinguish the brain patterns associated with the category of the foveated images. The performance of the classifier was above chance in the −135 – −105 ms interval and the −72 – 250 ms interval (p < 0.05, cluster permutations approach). The category classification gradually built up until peaking to 0.74 at 145 ms, after which they started to decrease (**Figure 3A**, red curve). These results demonstrate that foveal images are processed in succession at the feature and category level during natural viewing. Note that the classification performance at the category level began before the fixation onset on the foveal image (−135 – −105; −72 – 250 ms), providing evidence in favour of a parafoveal previewing at the category level; however, the classification accuracy increased dramatically ∼60 ms and ∼130 ms after fixation for respectively feature and category decoding.

**Figure 3.**
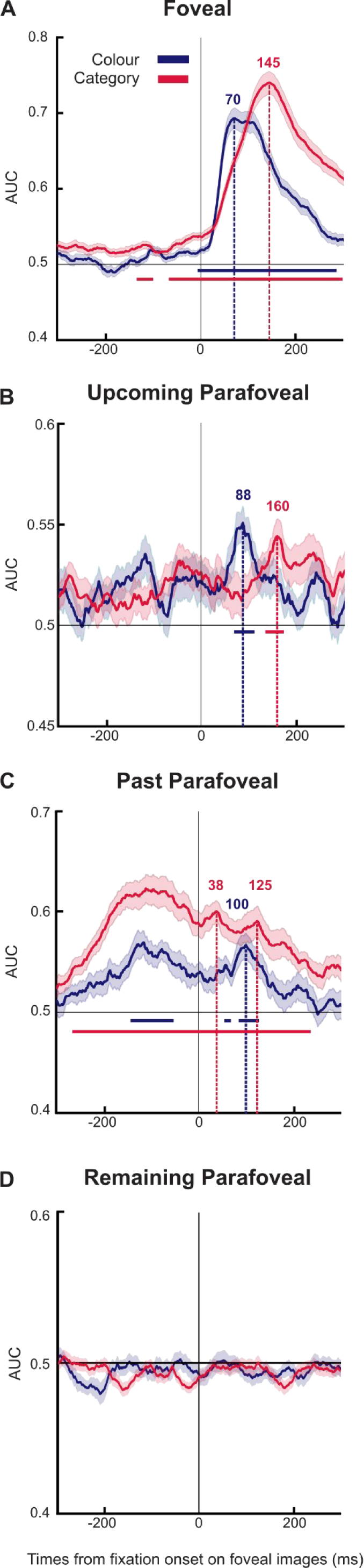
Decoding foveal, upcoming parafoveal, and past parafoveal images at the feature and category level. The feature and the category of foveal, upcoming parafoveal, and past parafoveal images can be identified from the MEG data during the foveation as shown by the classification analysis across time. The MEG data was aligned to fixation onset on foveal images (5-fold cross-validation; [−250; +250] ms; Area Under the Curve, AUC). Classification for colour (*greyscale vs colour*: blue line), and category (*animal vs food vs object*: red line), for **A.** Foveal images (currently fixated images), **B.** Upcoming parafoveal images (viewed after the current image), **C.** Past parafoveal images (viewed before the fixation on the current image). **D**. Remaining parafoveal image (not viewed after, not viewed before the current image). The shaded areas reflect the Standard Error of the Mean (SEM). A cluster permutation approach was used to identify significant time points (coloured horizontal lines).

### Decoding of parafoveal upcoming images at the feature and category level

Next, we investigated the parafoveal previewing at the feature and category level. We trained and tested the same classifier at each time point for the data aligned to the onset of the foveal images, to test whether we can classify the colour and category of the upcoming parafoveal images. As shown in **Figure 3B** (blue curve), the classifier could distinguish the colour of the parafoveal images, well above chance in the 68 – 105 ms interval (p < 0.05, cluster permutations approach). The decoding peaked to 0.55 at 88 ms. Similarly, the classifier was able to differentiate the category of the parafoveal images. The performance was above chance in the 132 – 172 ms interval (p < 0.05, cluster permutations approach), with a peak at 0.54 at 160 ms (**Figure 3B**, red curve). We here also observed a gradual increase and decrease of the performance around the peak, for both the colour and feature decoding, but less strongly than for the foveal classification. Our analysis shows that during unconstrained visual exploration, there is parafoveal previewing of upcoming saccade targets at both the feature and the category levels.

### Decoding of parafoveal past images at the feature and the category level

We then tested whether the feature and category of the image previously seen could be decoded. The classifier was trained and tested on data aligned according to fixation onset on the foveal images, to test whether we could classify the colour and the category of the foveal images viewed just before the saccade. As seen in **Figure 3C** (blue curve), we found that the classifier was able to categorize the colour of the parafoveal past images in the −150 – −60 ms interval (p < 0.05, cluster permutations approach). This is unsurprising as this is likely to reflect the image being on the fovea as t < 0 ms. More surprisingly, we found robust decoding of the past parafoveal image-colour during in the 54 – 62 ms interval following fixation, as well as in the 84 – 122 ms interval. The classification peaked to 0.57 at 100 ms. The classifier could also identify the category of the past parafoveal images from in the −270 –235 ms interval (p < 0.05, cluster permutations approach; **Figure 3C**, red curve), with two peaks during the foveation, an early peak to 0.6 at 38 ms, and a late peak to 0.59 at 125 ms. These results demonstrate that the neuronal activity reflecting both the feature and the category information of a given image is sustained after the subsequent saccade.

### Absence of decoding accuracy for parafoveal non-targets

We additionally computed classification accuracy for parafoveal images that were not targets of upcoming saccades. The classifier was trained and tested on data aligned according to fixation onset on the foveal images. As seen in **Figure 3D**, the classifier was not able to distinguish the colour (blue curve) and the category (red curve) of parafoveal images (AUC at the chance level). These results demonstrate that parafoveal images can only be decoded if they are included in the saccade goal, a notion consistent with pre-saccadic attention.

### Temporal generalization

To quantify the temporal generalization, the classifier was trained on all time points and tested on all time points, for colour and category classification. The temporal generalization analysis results in 2D matrices of classification performance, with the diagonal corresponding to when the classifier was trained and tested at the same time points (i.e., as in **Figure 3**). The classification rate above chance at the off-diagonal reflects that the training for these time points enables the classifier to generalize to other time points (**Figure 4**). This approach allows us to investigate how stable the neural code is across time [40]. For the colour classification, a succinct diagonal pattern was observed only in the foveal condition (**Figure 4A**) (p < 0.05, cluster permutations approach). In the upcoming parafoveal (**Figure 4C**) and the past parafoveal (**Figure 4E**) conditions, the classification of colour did not survive the multiple comparisons correction. These results suggest that the colour of natural images was processed transiently by the brain. For the classification of the images’ category, a square-like generalization matrix was observed off the diagonal in the foveal condition (**Figure 4B**), in the upcoming parafoveal condition (**Figure 4D**), and in the past parafoveal condition (**Figure 4F**) (p < 0.05, cluster permutations approach). This pattern suggests that the images’ category was encoded by a stable ensemble of neurons across time, for hundreds of milliseconds. In sum, although the neuronal pattern that encodes the colour of natural images was transient, the pattern encoding the category was stable across time. While the spatial brain patterns associated with the category classification seemed to be consistent in the foveal, upcoming parafoveal, and past parafoveal processes, the classification of colour appeared to be weaker and transient for the parafoveal images.

**Figure 4.**
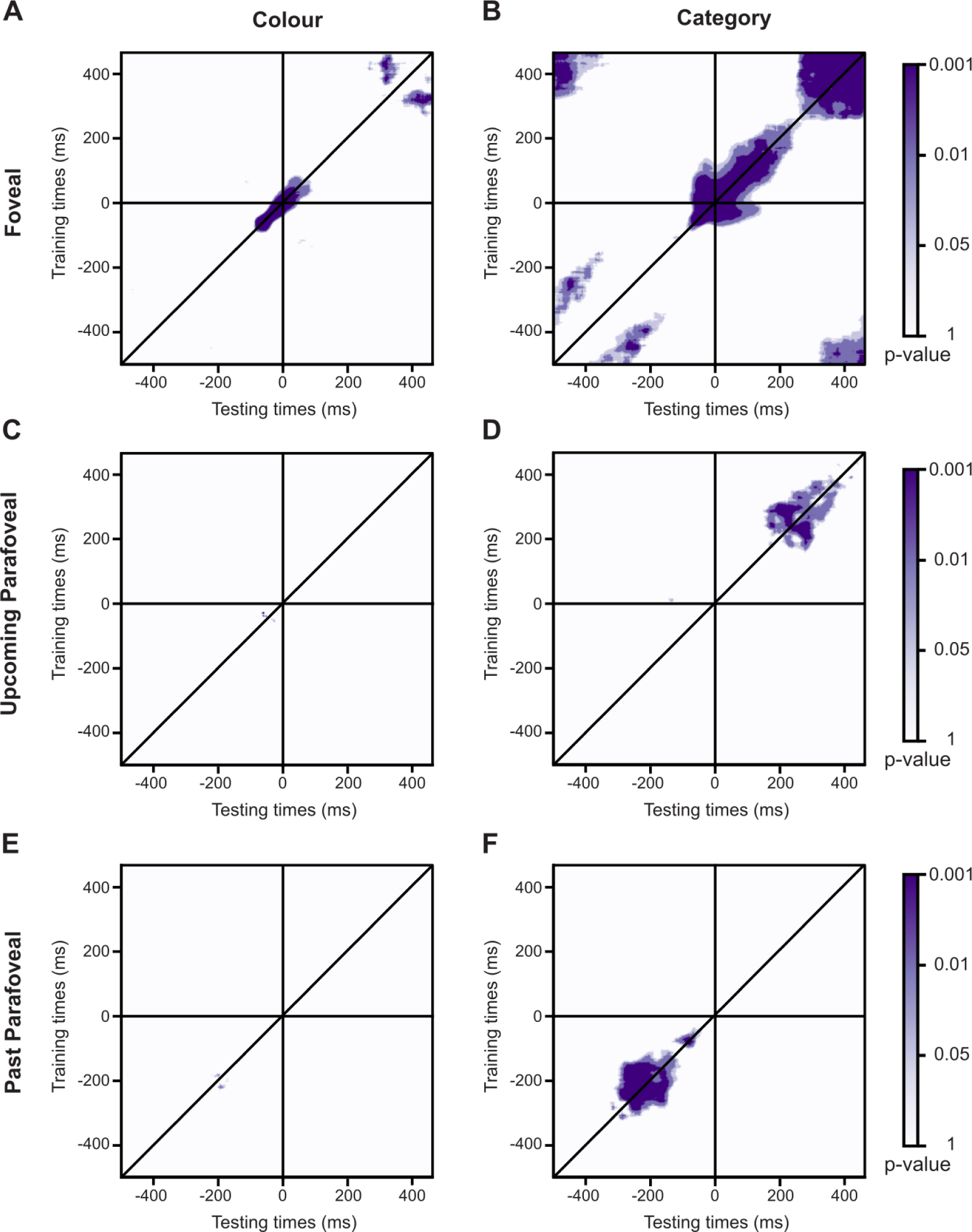
The decoding for colour classification was transient while stable across an ∼200 ms interval for category classification. Temporal generalization analysis on MEG data aligned to fixation onset on the foveal images **A.** Colour classification, Foveal images, **B.** Category classification, Foveal images, **C.** Colour classification, Upcoming parafoveal images, **D.** Category classification, Upcoming parafoveal images, **E.** Colour classification, Past parafoveal images, **F.** Category classification, Past parafoveal images. The colour scale reflects p-value from the cluster permutations approach, used to identify the significant time points with correction for multiple comparisons. A succinct diagonal pattern was observed in the temporal generalization of colour classification, while a square generalization matrix was noticed for the category classification. These results suggest that colour processing was transient, while category processing was stable for ∼200 across time.

### Sequential processing of foveal and parafoveal category information

Whether the processing of the foveal and parafoveal information is sequential or parallel along the visual hierarchy is debated. To address this question, we extracted the latency of the peaks of classification for each participant during the foveation, for the foveal and upcoming parafoveal images, with respect to colour and category classification (see **Figure 3A and B**). We used a Jackknife procedure to compare the latencies between the foveal and the parafoveal classification [41]. The latency differences were recomputed by sequentially removing a single participant from the sample of N=32 participants. The Jackknife statistics was then estimated by dividing the latency differences computed in the overall sample by the estimate of the standard error obtained in the subsamples. The Jackknife statistics was finally converted into a p-value. For the colour classification, we did not find a significant difference in the peak’s latencies between the foveal and the upcoming parafoveal conditions (p = 0.65, Cohen’s d = 0.23). Peak latencies across participants (n = 32) were 86 ± 30 ms (mean ± std) for the foveal image and 122 ± 71 ms for the parafoveal image (**Figure 5A**). For category classification, there was a significant difference in latencies between the foveal and the upcoming parafoveal images (p < 0.001, Cohen’s d = 1.43). Peak latency averaged across participants (n = 32) was (mean ± std) 139 ± 20 ms for the foveal images and 149 ± 75 ms for the upcoming parafoveal condition (**Figure 5B**). These findings might suggest sequential processing of the category information, delayed in terms of peak-times by 10 ms between the foveal and the parafoveal images. However, we also note that there was considerable overlap in the intervals of significant classifications between foveal and parafoveal images.

**Figure 5.**
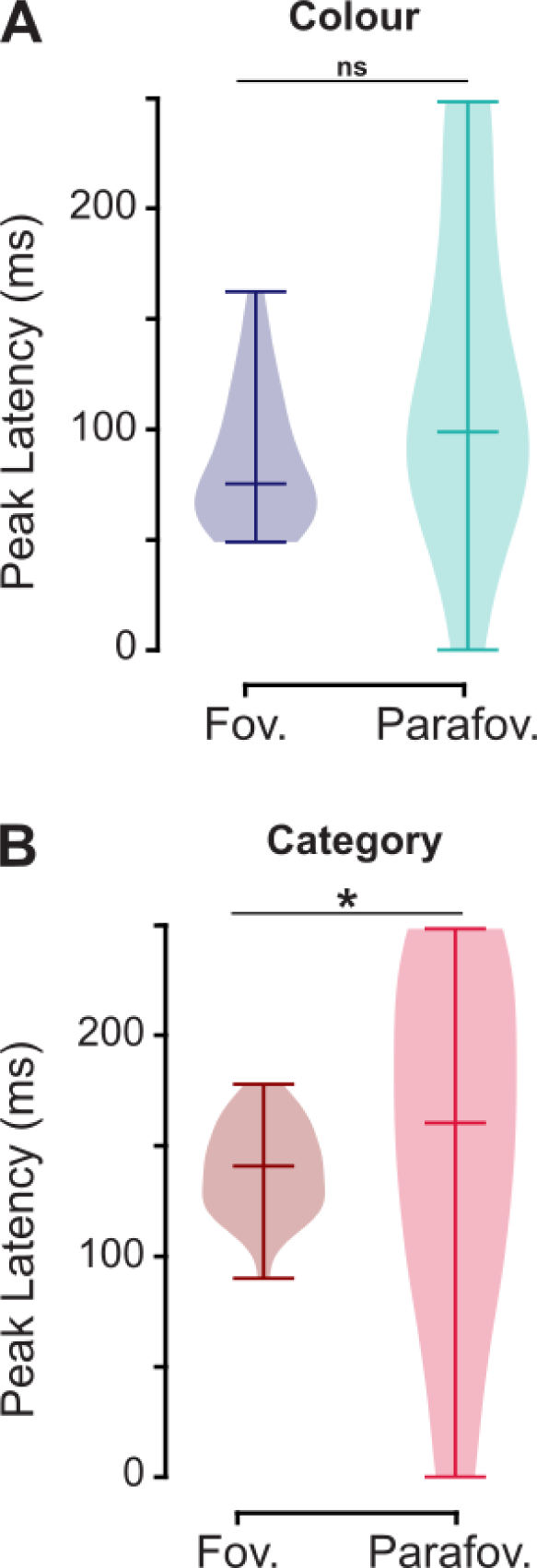
Category decoding peaked later for upcoming parafoveal compared to foveal images. The peak latencies computed during the foveation for the N=32 participants. **A.** Colour classification. No difference was observed for the colour categorization (Jackknife statistics, p = 0.65, Cohen’s d = 0.23). **B.** Category classification. The peak of classification was significantly later for the parafoveal compared to the foveal images for the object categorization (Jackknife statistics, p < 0.001, Cohen’s d = 1.43). The horizontal bar in the violin plots indicates the mean value.

### The decoding of features and the categories of foveal images predicted the behavioural performance

The decoding performance reflects how well the brain patterns associated with different conditions can be distinguished, but do they reflect the perception and maintenance of the objects? We hypothesized that the decoding would predict the behavioural performance. To test this hypothesis, the classifier was trained and tested on correct and incorrect trials separately, on foveal and upcoming parafoveal images. For foveal images, the classifier was able to categorize the colour in the 10 – 250 ms interval and in the 30 – 185 ms interval, for correct (light blue curve) and incorrect (dark blue curve) trials respectively (p < 0.05, cluster permutations approach; **Figure 6A**). A significant difference in the classification of features was observed between correct and incorrect trials in the 40 – 185 ms interval (we compared correct and incorrect conditions when their performance was above the chance level; p < 0.05, cluster permutations approach; **Figure 6A**). The classifier identified the category of foveal images, in the −25 – 250 ms interval and in the 45 – 250 ms interval, for correct (light red curve) and incorrect (dark red curve) trials respectively (p < 0.05, cluster permutations approach; **Figure 6B**). We found a significant difference in classification performance between correct and incorrect trials in the 28 – 250 ms interval (correct and incorrect conditions were compared when their performance was above the chance level; p < 0.05, cluster permutations approach; **Figure 6B**). Regarding the upcoming parafoveal images, the classifier was not able to identify either the colour or the category for correct and incorrect trials (performance was not significantly different from the chance level; p > 0.05, cluster permutations approach), likely because the number of trials was too low. Consequently, we cannot test whether the classification performance differs between correct and incorrect trials for upcoming parafoveal images. In sum, we found that decoding of the foveal images predicted behavioural performance.

**Figure 6.**
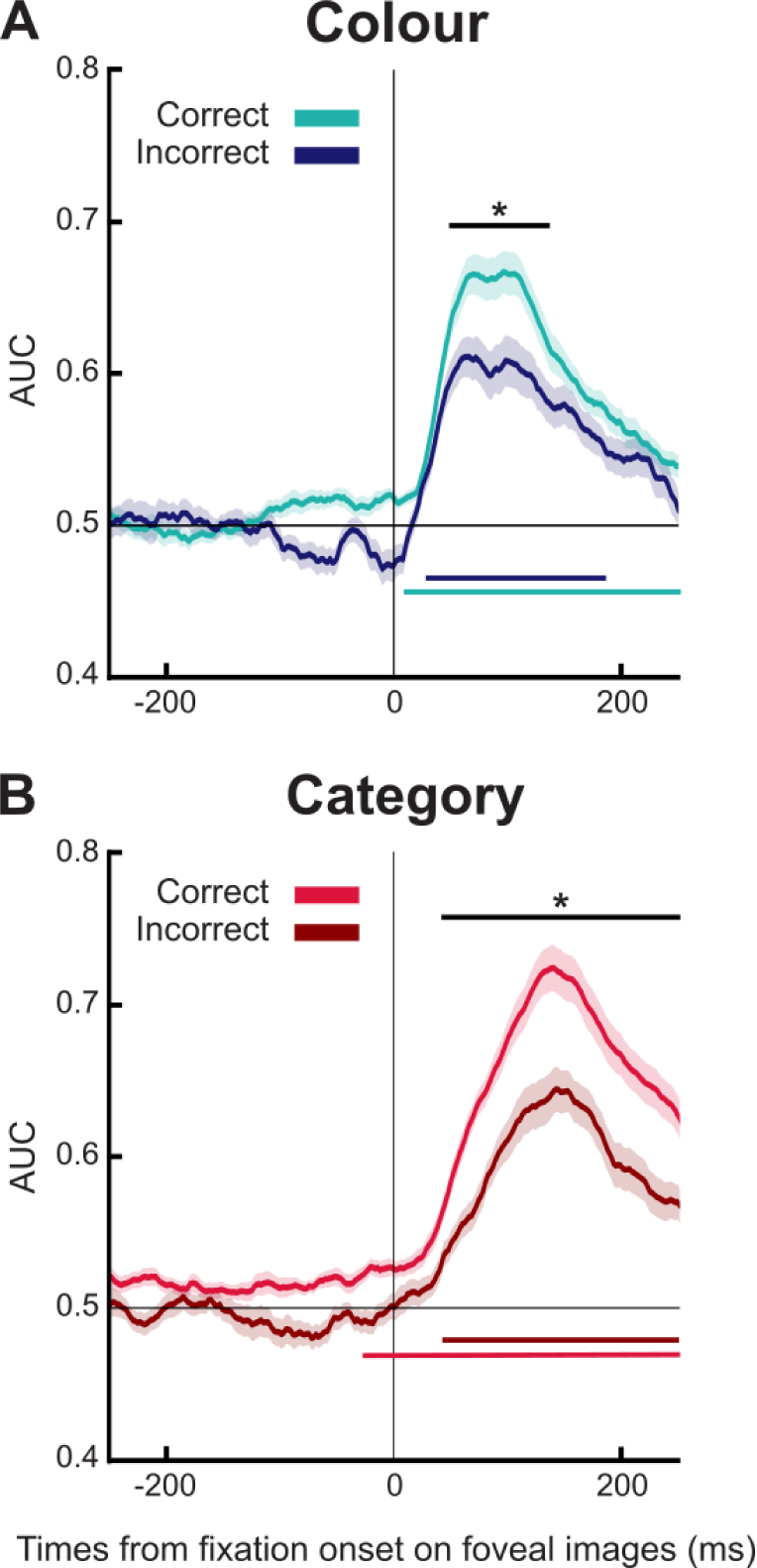
Feature and category decoding of foveal images higher for correct compared to incorrect trials. **A.** The feature (*greyscale vs colour*: blue lines) and **B.** The category (e.g. *animal vs food*: red lines) of foveal images are better identified for correct than incorrect trials in MEG data aligned to fixation onset on foveal images (5-fold cross validation; [−250; +250] ms; Area Under the Curve, AUC). The shaded areas reflect Standard Error of the Mean (SEM). A cluster permutations approach (p < 0.05) was used to identify the significant time points above the chance level (coloured horizontal lines), and to compare correct and incorrect trial classifications (black horizontal line) across time.

## Discussion

The purpose of this study was to investigate pre-saccadic attention, by uncovering how quickly and to what detail the brain processes visual information when we explore visual images. We applied multivariate pattern analysis (MVPA) to analyse MEG and eye-tracking data of participants viewing sets of seven natural images. These images belonged to different categories and were presented in greyscale or colour. The MVPA results showed that the brain processes images both in the fovea and the parafovea at the feature (greyscale vs colour) and category (e.g., food versus animal) levels. The parafoveal feature and category-specific decoding peaked at ∼90 ms and ∼160 ms respectively, whereas decoding of fixated objects peaked at ∼70 ms and ∼145 ms respectively. However, in support of parafoveal processing the category-decoding of fixated objects was robust already slightly before fixation. The magnitude of the classification of the foveated object predicted perceptual performance. Additionally, we found that feature and category-specific information about previously fixated objects in the parafoveal area was maintained for ∼ 230 ms after the eyes had moved away from them. Interestingly, the feature and category of non-target parafoveal images could not be decoded, suggesting that visual processing was constrained by the saccade goal. Overall, these findings showed that the brain can simultaneously extract detailed information about both foveal and parafoveal objects within an intersaccadic interval, and this may help to plan the next eye movement as well as to construct a representation of the full scene.

We found that the classification of upcoming parafoveal visual objects peaked at the feature and the semantic level at ∼90 ms and ∼160 ms respectively. This is consistent with a previous study using EEG and a paradigm involving controlled eye movements in which it was found that the category classification of parafoveal objects peaked at ∼160 ms [28]. We complement these findings by identifying a similar time course during unconstrained visual exploration. Another EEG study used free visual exploration and fixation-related potentials to show that the congruency of parafoveal visual target objects with respect to the scene was reflected in the N400 response before saccading to the parafoveal object [10]. We here extend these findings by demonstrating that the category-specific parafoveal processing builds up and peaks already at ∼160 ms after fixating on the pre-target object. Importantly, the parafoveal processing of the feature and the category was specific to objects of the next saccade goal. Considering that the oculomotor system takes 100 ms to initiate and execute a saccade [3], and that humans typically saccades every ∼250-300 ms [36], our results provide electrophysiological evidence that semantic information of the parafoveal object can be extracted sufficiently fast (within ∼150 ms) to impact saccade plans in natural viewing.

In our study, the classification of feature and the semantic information of foveal objects peaked at 70 ms and 145 ms respectively, after the fixation onset. Thus, the parafoveal preview resulted in only a modest if any speed up in the categorization of visual objects after fixations, given that other studies have found a peak in classification at ∼150 ms without previewing [42–44]. These considerations only apply to the peak of classification. As we showed in **Figure 3A**, the object category could be decoded above chance already prior to the saccade, from −135 to −105 ms, and from −72 ms to fixation onset. This provides evidence for semantic evidence starting to accumulate of parafoveal objects already prior to the saccade. In our design, we did not have a condition preventing parafoveal processing. Other studies using tasks with controlled eye movements [17–25, 28] have found evidence for preview benefit, where some suggest that parafoveal previewing reduces the subsequent processing time by at least 30 ms [15, 28]. In future studies, it would be of great interest to extend our design such that previewing is manipulated. This could be done using a gaze-contingent paradigm in which the parafoveal object is replaced (or not) when saccading towards it. Our multivariate approach could then be used to compare the classification time after a preview versus a non-preview has been performed. This will allow for estimating the precise benefit of parafoveal processing in natural viewing conditions.

Our study revealed that feature decoding of foveal objects was robust in an 8 – 310 ms interval but peaked at 70 ms. Decoding the category of foveal objects was robust in the −135 – −105 ms and −72 – 250 ms intervals in relation to fixation but peaked at 145 ms. The peak-decoding of feature and category decoding of upcoming parafoveal objects were identified at 88 ms (robust in a 68 –105 ms interval) and 160 ms (robust in a 132 –172 ms interval) respectively. These results inform the debate regarding sequential versus parallel visual processing. First, our findings provide evidence against a strict serial mechanism, relying on parafoveal processing starting once the foveal object has been fully identified [45, 46]. We base this argument on feature-processing of the parafoveal visual object (peak at 88 ms, onset at 68 ms) starting already before the category-specific processing of the foveal object (peak at 145 ms, onset before fixation). In regard to mechanisms emphasising parallel processing [47], we did find considerable overlap in the time interval in which foveal and parafoveal objects were decoded both in terms of feature- and category-specific processing. As a challenge to a strict parallel processing scheme, our analysis revealed a significant ∼15-ms delay in the peak of decoding between the foveal and parafoveal processing of the category. While there was a difference (∼12 ms) in the peak of feature-specific processing, this was not statistically different. In addition, the colour and category processing of parafoveal objects started (onset at 68 and 132 ms respectively) later than for the foveal object (onset at 8 and −72 ms respectively). The onsets of feature and category decoding of parafoveal objects were in fact closer in time to the peak of foveal decoding (peak at 70 and 145 ms respectively). These differences in time suggested that parallel processing cannot fully account for the observed results. These results are closely in line with the idea of a pipelining mechanism [48]. According to this idea, several objects can be processed simultaneously but at different levels of the visual hierarchy. The sequential activation of the category-specific processing is aligned with the pipeline mechanism. In sum, while we provided evidence against the serial theory, our findings are better aligned with parallel or pipelining theories. Future studies would be necessary to further uncover the temporal dynamics and the associated mechanisms of foveal and parafoveal processing along the visual hierarchy.

We found that the classification performance at the feature and the category level for foveal images reflected the behavioural performance in the memory task. These findings suggest that the distributed brain responses are directly related to how well the images are perceived and retained. However, we were not able to link the decoding of parafoveal objects to performance. This may be a matter of low signal-to-ratio as there were not too many incorrect trials. In the future, increasing the number of trials as well as task difficulty would help to determine if the parafoveal processing also predicts behavioural and memory performance.

Finally, we found that the feature and the semantic information of past parafoveal images remained present until ∼230 ms after saccade onset. This finding is in line with studies based on controlled eye movements showing that brain data associated with low- and high-level visual features of past parafoveal objects remained present for 100-200 ms after the saccade [28, 34]. We here show that these findings generalize to free-viewing conditions. Our finding supports the notion that visual information can be transferred across saccades [29–31]. In future research, it would be interesting to uncover how information from past and current saccades contributes to building the full percept of a visual scene.

While the participants scanned the display at their own volition, the configuration of the 7 images was somewhat artificial. The next step could be to develop the approach to study the exploration of real natural scenes. The objects in the natural scenes as well as their location could be labelled using automatic tools derived from machine learning [49, 50]. If successful, a similar approach could be used for videos. Eventually the study could be conducted outside the laboratory where the brain activity is measured from participants walking in a natural environment with eye-tracking glasses and the brain activity measured using a mobile EEG.

Our study provides electrophysiological evidence that parafoveal images are processed at the feature and category levels during natural visual exploration, within ∼150 ms after fixation onset. This demonstrates that semantic information of parafoveal objects can be extracted sufficiently fast to guide saccades during natural viewing and potentially support the parafoveal preview benefit. Our results also demonstrate that within an inter-saccade interval, information about past, current, and upcoming objects is represented in the brain. This information might be part of the building blocks supporting stable perception in the presence of saccades.

## Materials and Methods

### Participants

36 participants (29 females, 1 left-handed, mean ± sd age: 21.4 ± 3 years) were included in the study. All participants had a corrected-to-normal vision and were free from medication affecting the central nervous system, reported no history of psychiatric or neurological disorders. The study was approved by the University of Birmingham Ethics Committee. All participants gave their written informed consent and received a monetary compensation or course credits for their participation.

### Stimuli

The stimuli used were fixation crosses, natural images, and masks. The fixation cross was black, with arms of 0.2 degrees of visual angles (°) of length and 0.05° of width. 1500 natural images of our three categories, *animal*, *food*, and *object*, were selected from the THINGS database [35]. The natural images were 3° x 3° presented in colour or grayscale. In each trial, one image was displayed in the centre on the screen, surrounded by 6 other images (at 0°, 60°, 120°, 180°, 240° and 300°), with a 1° distance between each other’s borders (**Figure 1A**). The proportion of image categories and grey vs colour scales was balanced within and between trials. The 3° x 3° masks were patches of random grey pixels.

### Procedure

Participants were seated comfortably in the MEG gantry, 145 cm from the projection screen in a dimly lit magnetically shielded room. Participants performed 10 blocks of 30 trials, with the presentation of 7 images per trial. In total, 2100 images were presented throughout the experiment, with approximatively 20% of repeated images. As shown in **Figure 1A**, each trial began when participants were successfully maintaining fixation, with the presentation of a fixation cross for 500 ms, presented on a grey background. Then 7 natural images were presented for 4000 ms. Participants were instructed to freely explore the 7 images and saccades were allowed. Then followed a masked presented for 2000 ms after which the 7 images were presented again, except for one image that was changed (a different image, belonging to the same category and colourscale as the initial one, e.g. a greyscale fox turning into a greyscale cat, **Figure 1A**). Participants had to identify which image was different from the initial presentation. They had no time-limit to respond. A number (1 to 7) was presented above each image, as well as a figure showing the link between the image numbers and the response buttons. The trials were separated by random intervals varying from 500 to 1000 ms (in 100 ms steps).

### Equipment

The experimental protocol was designed using the PsychToolbox 3.0.12, implemented in Matlab 2015b (The MathWorks, Natick, MA). Visual stimuli were displayed with a ProPixx Projector (VPixx Technologies, Saint-Bruno, QC, Canada), on a 71.5 by 40.2 cm projection screen (1920 by 1080 pixels; 120 Hz refresh rate).

#### MEG

MEG data were acquired using a 306-sensor TRIUX Elekta Neuromag system with 204 orthogonal planar gradiometers and 102 magnetometers (Elekta, Finland). Participants were then seated under the MEG gantry with the back rest at 60° angle. The data were band-pass filtered from 0.1 to 330 Hz and then sampled at 1000 Hz.

Prior to the MEG study, a Polhemus Fastrack electromagnetic digitizer system (Polhemus Inc., USA) was used to digitize the locations of three fiducial points (nasion, left and right peri-auricular points) and of four head position indicator coils (HPI coils). Two HPI coils were placed on the left end right mastoid bone, and the two others on the forehead with at least 3 cm distance. At least 300 extra points on the scalp were additionally digitalized.

#### Eye Tracker

An infrared video-camera system (EyeLink 1000 Plus, SR Research, Ottawa, Canada) was used to record participants’ eye movements sampled at 1000 Hz. A 9-point calibration was performed at the beginning of the experiment and one-point drift corrections were applied at the beginning of every block.

### Analyses

Analyses were performed with custom software written in Python 3.10.9 (Python Software Foundation. Python Language Reference. Available at http://www.python.org) and figures were plotted with *Matplotlib* library [51]. MEG analyses were performed using MNE 1.0.3 [52].

### Behavioural analysis

Behavioural performance was computed as the hit rate, i.e., percentage of correct responses, and median reaction times for correct and incorrect trials, for each participant.

### Eye data analysis

The following eye metrics were extracted from the EDF files provided by the EyeLink toolbox, during the initial presentation of the images: number of fixations per trial, fixation durations, number of saccades per trial, and saccade duration, for correct and incorrect trials.

### MEG analysis

#### Preprocessing

MEG data were preprocessed following the standards defined in the FLUX Pipeline [53]. Continuous head movements were estimated by computing the time-varying amplitude of the HPI coils. Sensors with no signal or excessive artefacts were removed using the *mne.preprocessing.find_bad_channels_maxwell* function, and a low-pass filter at 150 Hz was applied to remove the activity from the HPI coils. A Maxwell filter including spatiotemporal signal-space separation (tSSS) was applied to the MEG signal. This procedure removes low-frequency artifacts and performs head movement compensation. Muscle artefacts, defined as activity in the 110 – 140 Hz band exceeding a z-score threshold at 10, were annotated as artefactual. Trials with muscle-artifacts were further rejected. An ICA decomposition was performed on the data bandpass filtered at 1 – 40 Hz. The components were inspected visually for each participant to remove the ocular and cardiac artefacts’ activity in the unfiltered MEG data (typically 3 – 5 components in each participant).

#### Epoching

Data were extracted in −1 to +1 s epochs aligned to the fixation onset on the images displayed during the initial presentation (fixations on the background were not analysed). Epochs with a sensor activity exceeding a 5000 fT/cm threshold for the gradiometers, and 5000 fT for the magnetometers, were rejected. Epochs were then downsampled to 500 Hz. We discarded epochs with fixation durations below 80 ms and above 1000 ms. As explained in **Figure 1B**, the epochs were then labelled according to the colour (*colour* vs *grey*) and the category (*animal, food,* or *object*) of:

- parafoveal past images (fixated before the currently image),
- foveal images (currently fixated images),
- parafoveal upcoming images (fixated after the current image),
- one parafoveal remaining image (not fixated either before either after the current image).

The parafoveal visual field encompasses objects between 2° and 5° of eccentricity. Consequently, past images viewed before the current fixation in the periphery were not analysed. Similarly, upcoming images in the periphery relative to the current fixation were not analysed (**Figure 1B**). In addition, we did not consider parafoveal past images already visited during the trial, as well as for parafoveal upcoming images. Eventually, remaining images in the periphery relative to the current fixation were also not analysed.

#### Quality check of data

Fixation-Related Fields (FRFs) were computed for the foveal condition across all colour and object categories, for each participant, and plotted for an occipital sensor (MEG2032). Four participants who did not show the typical ERFs (P1m, N2m, P2m) were excluded from the analysis, and remaining 32 participants analysed in total.

#### Multivariate Pattern Analysis (MVPA)

Multivariate pattern analysis was applied to the MEG data to investigate whether the brain pattern associated with colour categorization (*grey* vs *colour*) and object categorization (*animal* vs *food* vs *object*) can be classified in relation to parafoveal past images, foveal images, and in parafoveal upcoming images (**Figure 1B**). For classification, we used a support vector machine [54] from *Scikit-learn* library [55], with a 5-fold cross-validation procedure. Epochs were cropped from −0.5 to +0.5 s including electrophysiological activity from both the gradiometers and the magnetometers (306 sensors). For each time point, we considered the data in a 50-ms time window (25 samples centred around the time point), resulting in a feature vector with 25 x N-sensor samples (also termed time-delay embedding; code available at [56]). It also permits a greater resilience to the varying activation delays across participants. To further increase the signal-to-noise ratio, we averaged 10 trials (randomly selected) in the training set and in the testing test to create so-called *super-trials* for each category [57, 58]. The creation of super-trials and the classification were repeated 10 times, and the final classification performance was obtained by averaging the classification rates across the 10 repetitions. The classification rate was reported as Area Under the Curve (AUC).

The MVPA was applied to the parafoveal past images, the foveal images, the parafoveal upcoming images, and the parafoveal remaining images. We investigated whether the classifier could disentangle greyscale vs colour images, and the category. Note that for the category, we averaged the performance over the three classifications: *animal* vs *food, animal* vs *object,* and *food* vs *object*. The MVPA was computed for each participant, and the classification performance were averaged across participants.

In addition, a MVPA with generalization across time was performed [40] (except for parafoveal remaining images regarding the results of the classic MVPA). The classifier was trained at a given time-point and tested on every time points, resulting in a 2D-matrix of classification rate, with the diagonal corresponding to when the trained time point matches the testing time point (i.e., MVPA across time).

To investigate whether the classification performance was modulated by the behavioural performance, the MVPA was further applied to the foveal images and the parafoveal upcoming images, separately for correct and incorrect trials, for the colour and the category. A subsampling procedure was added to the *super-trials* generation to have an equal number of correct and incorrect trials per participant.

### Statistical analysis

Behavioural and eye data were compared between correct and incorrect trials with t-test from the *Pingouin* library [59]. To investigate at which time points the classifier performance was above the chance level (0.5), we used a cluster-permutation approach to control for multiple comparisons over time points [37–39]. For ease of computation, the chance level was set at 0. For each of the 1,500 repetitions, the classification performance was randomly multiplied by 1 or −1 with equivalence across participants, and an independent t-test was computed against zero (chance level) with *Scipy* library [60], across all time points. The maximum t-value was considered at each repetition, leading to a distribution of t-values, from which we extracted a threshold t-value for alpha = 5%. A t-value above the threshold t-value was considered significant.

A similar cluster-permutation approach was used to test at which time points the classifier performance differs between correct and incorrect trials. For each of the 1,500 repetitions, the classification performance was randomly assigned to the correct or incorrect conditions across participants, and an independent t-test was computed across all time points. The maximum t-value was considered at each repetition to obtain a distribution of t-values. A t-value above a 5% threshold was considered significant.

To evaluate the latency of the classification peaks, we identified the time-points of peaking in each participant in a 0 – 250 ms interval according to fixation onset for the different conditions. These latencies were compared between the foveal and the parafoveal conditions by applying a Jackknife procedure (also called leave-one-out; [41]). In our application of the Jackknife procedure [41], the difference in latencies was computed from a subsample including all participants except one. The procedure was repeated by removing data from each participant sequentially, resulting in N subsample differences for a sample size of N participants. Then the Jackknife estimate of the standard error of the difference is:

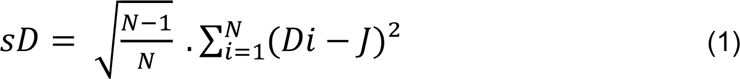

with N the number of participants, Di the subsample difference for each participant exclusion, and J the mean of the differences obtained in the subsamples. The Jackknife statistics was then computed with the following formula:

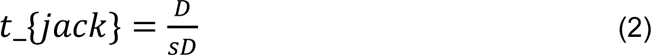

with D the difference in latencies observed in the overall sample, and sD the estimate of its standard error obtained from Equation 1. According to the null hypothesis, the Jackknife statistics t_{jack} should have approximatively a t distribution with N – 1 degrees of freedom because D is approximatively normal with a mean of zero and sD is an estimate of the standard deviation D. Thus, the Jackknife statistics can be converted to a standard p-value with the following formula:

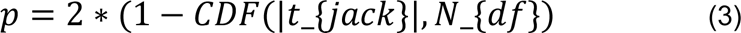

with CDF the cumulative distribution function, t_{jack} the Jackknife statistics, and N_{df} the degrees of freedom.

## Coda and Data sharing

After publication, data will be available on a Birmingham University server, and code will be available at https://github.com/CamilleFakche.

## Bibliography

1. Yarbus AL. Eye movements and vision. Springer. 1967.

2. Prasad S, Galetta SL. Anatomy and physiology of the afferent visual system. Handbook of clinical neurology. 2011; 102, 3–19.

3. Otero-Millan J, Troncoso XG, Macknik SL, Serrano-Pedraza I, Martinez-Conde S. Saccades and microsaccades during visual fixation, exploration, and search: foundations for a common saccadic generator. Journal of vision. 2008; 8(14), 21–21.

4. Henderson JM, Weeks Jr PA, Hollingworth A. The effects of semantic consistency on eye movements during complex scene viewing. Journal of experimental psychology: Human perception and performance. 1999; 25(1), 210.

5. Bonitz VS, Gordon RD. Attention to smoking-related and incongruous objects during scene viewing. Acta psychologica. 2008; 129(2), 255–263.

6. Borges MT, Fernandes EG, Coco MI. Age-related differences during visual search: the role of contextual expectations and cognitive control mechanisms. Aging, Neuropsychology, and Cognition. 2020; 27(4), 489–516.

7. LaPointe MR, Milliken B. Semantically incongruent objects attract eye gaze when viewing scenes for change. Visual Cognition. 2016; 24(1), 63–77.

8. Nuthmann A, De Groo F, Huettig F, Olivers CN. Extrafoveal attentional capture by object semantics. PLoS One. 2019; 14(5), e0217051.

9. Cimminella F, Sala SD, Coco MI. Extra-foveal processing of object semantics guides early overt attention during visual search. Attention, Perception, & Psychophysics. 2020; 82(2), 655–670.

10. Coco MI, Nuthmann A, Dimigen O. Fixation-related brain potentials during semantic integration of object–scene information. Journal of Cognitive Neuroscience. 2020; 32(4), 571–589.

11. Rayner K. Eye movements in reading and information processing: 20 years of research. Psychological bulletin. 1998; 124(3), 372.

12. Huber-Huber C, Buonocore A, Melcher D. The extrafoveal preview paradigm as a measure of predictive, active sampling in visual perception. Journal of vision. 2021; 21(7), 12–12.

13. Henderson JM, Pollatsek A, Rayner K. Effects of foveal priming and extrafoveal preview on object identification. Journal of Experimental Psychology: Human Perception and Performance. 1987; 13(3), 449.

14. Henderson JM. Identifying objects across saccades: effects of extrafoveal preview and flanker object context. Journal of Experimental Psychology: Learning, Memory, and Cognition. 1992; 18(3), 521.

15. Pollatsek A, Rayner K, Henderson JM. Role of spatial location in integration of pictorial information across saccades. Journal of Experimental Psychology: Human Perception and Performance. 1990; 16(1), 199.

16. Henderson JM, Pollatsek A, Rayner K. Covert visual attention and extrafoveal information use during object identification. Perception & psychophysics. 1989; 45(3), 196–208.

17. Morgan JL, Meyer AS. Processing of extrafoveal objects during multiple-object naming. Journal of Experimental Psychology: Learning, Memory, and Cognition. 2005; 31(3), 428.

18. Schotter ER, Ferreira VS, Rayner K. Parallel object activation and attentional gating of information: Evidence from eye movements in the multiple object naming paradigm. Journal of Experimental Psychology: Learning, Memory, and Cognition. 2013; 39(2), 365.

19. Ganmor E, Landy MS, Simoncelli EP. Near-optimal integration of orientation information across saccades. Journal of vision. 2015; 15(16), 8–8.

20. Wijdenes LO, Marshall L, Bays PM. Evidence for optimal integration of visual feature representations across saccades. Journal of Neuroscience. 2015; 35(28), 10146–10153.

21. Castelhano MS, Pereira EJ. The influence of scene context on parafoveal processing of objects. Quarterly Journal of Experimental Psychology. 2018; 71(1), 229–240.

22. Stewart EE, Schütz AC. Optimal trans-saccadic integration relies on visual working memory. Vision Research. 2018; 153, 70–81.

23. Kong G, Kroell LM, Schneegans S, Aagten-Murphy D, Bays PM. Transsaccadic integration relies on a limited memory resource. Journal of Vision. 2021; 21(5), 24–24.

24. Huber-Huber C, Buonocore A, Dimigen O, Hickey C, Melcher D. The peripheral preview effect with faces: Combined EEG and eye-tracking suggests multiple stages of trans-saccadic predictive and non-predictive processing. NeuroImage. 2019; 200, 344–362.

25. Buonocore A, Dimigen O, Melcher D Post-saccadic face processing is modulated by pre-saccadic preview: Evidence from fixation-related potentials. Journal of Neuroscience. 2020; 40(11), 2305–2313.

26. Ehinger BV, König P, Ossandón JP. Predictions of visual content across eye movements and their modulation by inferred information. Journal of Neuroscience. 2015; 35(19), 7403–7413.

27. De Lissa P, McArthur G, Hawelka S, Palermo R, Mahajan Y, Degno F, Hutzler F. Peripheral preview abolishes N170 face-sensitivity at fixation: Using fixation-related potentials to investigate dynamic face processing. Visual Cognition. 2019; 27(9-10), 740–759.

28. Edwards G, VanRullen R, Cavanagh P. Decoding trans-saccadic memory. Journal of Neuroscience. 2018; 38(5), 1114–1123.

29. Melcher D, Colby CL. Trans-saccadic perception. Trends in cognitive sciences. 2008; 12(12), 466–473.

30. Cavanagh P, Hunt AR, Afraz A, Rolfs M. Visual stability based on remapping of attention pointers. Trends in cognitive sciences. 2010; 14(4), 147–153.

31. Herwig A. Transsaccadic integration and perceptual continuity. Journal of vision. 2015; 15(16), 7–7.

32. Wolf C, Schütz AC. Trans-saccadic integration of peripheral and foveal feature information is close to optimal. Journal of vision. 2015; 15(16), 1–1.

33. Paeye C, Collins T, Cavanagh P Transsaccadic perceptual fusion. Journal of Vision. 2017; 17(1), 14–14.

34. Fabius JH, Fracasso A, Acunzo DJ, Van der Stigchel S, Melcher D. Low-level visual information is maintained across saccades, allowing for a postsaccadic handoff between visual areas. Journal of Neuroscience. 2020; 40(49), 9476–9486.

35. Hebart MN, Dickter AH, Kidder A, Kwok WY, Corriveau A, Van Wicklin C, Baker CI. THINGS: A database of 1,854 object concepts and more than 26,000 naturalistic object images. PloS one. 2019; 14(10), e0223792.

36. Skaramagkas V, Giannakakis G, Ktistakis E, Manousos D, Karatzanis I, Tachos NS, et al. Review of eye tracking metrics involved in emotional and cognitive processes. IEEE Reviews in Biomedical Engineering. 2021; 16, 260–277.

37. Nichols TE, Holmes AP. Nonparametric permutation tests for functional neuroimaging: a primer with examples. Human brain mapping. 2002; 15(1), 1–25.

38. Maris E. Statistical testing in electrophysiological studies. Psychophysiology. 2012; 49(4), 549–565.

39. Winkler AM, Ridgway GR, Webster MA, Smith SM, Nichols TE. Permutation inference for the general linear model. Neuroimage. 2014; 92, 381–397.

40. King JR, Dehaene S. Characterizing the dynamics of mental representations: the temporal generalization method. Trends in cognitive sciences. 2014; 18(4), 203–210.

41. Miller J, Patterson TUI, Ulrich R. Jackknife-based method for measuring LRP onset latency differences. Psychophysiology. 1998; 35(1), 99–115.

42. Hung CP, Kreiman G, Poggio T, DiCarlo JJ. Fast readout of object identity from macaque inferior temporal cortex. Science. 2005; 310(5749), 863-866.

43. Cichy RM, Pantazis D, Oliva A. Resolving human object recognition in space and time. Nature neuroscience. 2014; 17(3), 455–462.

44. Bezsudnova Y, Quinn AJ, Wynn SC, Jensen O. Spatiotemporal properties of common semantic categories for words and pictures. Journal of Cognitive Neuroscience. Forthcoming.

45. White AL, Boynton GM, Yeatman JD. You can’t recognize two words simultaneously. Trends in Cognitive Sciences. 2019; 23(10), 812–814.

46. Reichle ED, Liversedge SP, Pollatsek A, Rayner K. Encoding multiple words simultaneously in reading is implausible. Trends in cognitive sciences. 2009; 13(3), 115–119.

47. Snell J, Grainger J. Readers are parallel processors. Trends in Cognitive Sciences. 2019; 23(7), 537–546.

48. Jensen O, Pan Y, Frisson S, Wang L. An oscillatory pipelining mechanism supporting previewing during visual exploration and reading. Trends in cognitive sciences. 2021; 25(12), 1033–1044.

49. Ramík DM, Sabourin C, Moreno R, Madani K. A machine learning based intelligent vision system for autonomous object detection and recognition. Applied intelligence. 2014; 40, 358–375.

50. Wäldchen J, Mäder P. Machine learning for image based species identification. Methods in Ecology and Evolution. 2018; 9(11), 2216–2225.

51. Hunter JD. Matplotlib: A 2D graphics environment. Computing in science & engineering. 2007; 9(03), 90–95.

52. Gramfort A, Luessi M, Larson E, Engemann DA, Strohmeier D, Brodbeck C, et al. MEG and EEG data analysis with MNE-Python. Frontiers in neuroscience. 2013; 7, 70133.

53. Ferrante O, Liu L, Minarik T, Gorska U, Ghafari T, Luo H, Jensen O. FLUX: A pipeline for MEG analysis. NeuroImage. 2022; 253, 119047.

54. Cortes C, Vapnik V. Support-vector networks. Machine learning. 1995; 20, 273–297.

55. Pedregosa F, Varoquaux G, Gramfort A, Michel V, Thirion B, Grisel O, et al. Scikit-learn: Machine learning in Python. the Journal of machine Learning research. 2011; 12, 2825–2830.

56. Rohrbacker N. Analysis of electroencephalogram data using time-delay embeddings to reconstruct phase space. Dynamics at the Horsetooth. 2009; 1, 1–11.

57. Isik L, Meyers EM, Leibo JZ, Poggio T. The dynamics of invariant object recognition in the human visual system. Journal of neurophysiology. 2014; 111(1), 91–102.

58. Grootswagers T, Wardle SG, Carlson TA. Decoding dynamic brain patterns from evoked responses: A tutorial on multivariate pattern analysis applied to time series neuroimaging data. Journal of cognitive neuroscience. 2017; 29(4), 677–697.

59. Vallat R. Pingouin: statistics in Python. J. Open Source Software. 2018; 3(31), 1026.

60. Virtanen P, Gommers R, Oliphant TE, Haberland M, Reddy T, Cournapeau D, et al. SciPy 1.0: fundamental algorithms for scientific computing in Python. Nature methods. 2020; 17(3), 261–272.

